# *Impatiens banen* sp. nov. & *I. etugei* sp. nov. (Balsaminaceae), threatened lowland species of the Cross-Sanaga Interval, Cameroon

**DOI:** 10.1101/2022.05.01.490166

**Authors:** Martin Cheek, Iain Darbyshire, Jean Michel Onana

## Abstract

We describe two range-restricted new species to science from the genus *Impatiens* (Balsaminaceae), both threatened, from the Cross-Sanaga Interval of western Cameroon. The first, *Impatiens banen*, appears to be restricted to an open seepage microhabitat on granitic inselbergs in the lowland-submontane forest zone of the Ebo Forest in Littoral Region and is provisionally assessed using the 2012 IUCN standard as Vulnerable. Sharing characters with *Impatiens burtonii* and *I. mannii*, it differs from both, and appears to be unique in Cameroon *inter alia* 1) in the bicolored united lateral petals, the upper petals being white, while the lower petals are an intense pink-purple, 2) the hairy, filamentous spur, purple with a white apex, is curved along its length (through nearly 360°), almost describing a circle. Inselberg-specific species are unusual in *Impatiens*.

The second species, *Impatiens etugei*, of the *I. macroptera* aggregate, is restricted to rocks in the Mutel River of the Kom Wum Forest Reserve of NorthWest Region and is assessed as Critically Endangered. Having similarities with *I. mackeyana* and *I. letouzeyi*, it differs from other species in the aggregate *inter alia* by having opposite leaves (vs always alternate), flower exterior white (vs pink or pink-purple), and in the dorsal petal having a pair of lateral projections (vs projections absent).

## Introduction

As part of the project to designate Important Plant Areas (IPAs) in Cameroon (also known as Tropical Important Plant Areas or TIPAs, https://www.kew.org/science/our-science/projects/tropical-important-plant-areas-cameroon), we are striving to name, assess the conservation status and include in TIPAs (Darbyshire *et al*. 2017) rare and threatened plant species in the threatened natural habitat of Cameroon.

New species to science from Cameroon are being published steadily, from herbs of waterfalls, forest shrubs to canopy trees (Achoundong *et al*. 2021; Alvarez-Aguirre *et al*. 2021; Cheek & Onana 2021; Cheek *et al*. 2021a; 2021b; in press; Gosline *et al*. 2022). Many of these species, including those described in this paper, were collected as part of a programme to produce a series of conservation checklists (see below) for areas of intact natural habitat ranging over much of the Cross-Sanaga interval (Cheek *et al*. 2001). The Cross-Sanaga has the highest vascular plant species, and highest generic diversity per degree square in tropical Africa (Barthlott *et al*. 1996; Dagallier *et al*. 2020, respectively), including endemic genera such as *Medusandra* Brenan (Peridiscaceae, Breteler *et al*. 2015; Soltis *et al*. 2007). However, natural habitat is being steadily cleared, predominantly for agriculture. Eight hundred and fifteen species of vascular plant were Red Listed at the global level for Cameroon (Onana & Cheek 2011), many of them confined to the Cross-Sanaga.

In this paper we describe two new species of *Impatiens* (Balsaminaceae), both apparently range-restricted and threatened. The first, *Impatiens banen* (Fig. 1 below) appears restricted to granitic inselbergs in the Ebo Forest of Littoral Region. The second, *Impatiens etugei*, to rocks in the river of the Kom Wum Forest Reserve of NW Region.

**Fig. 1.**
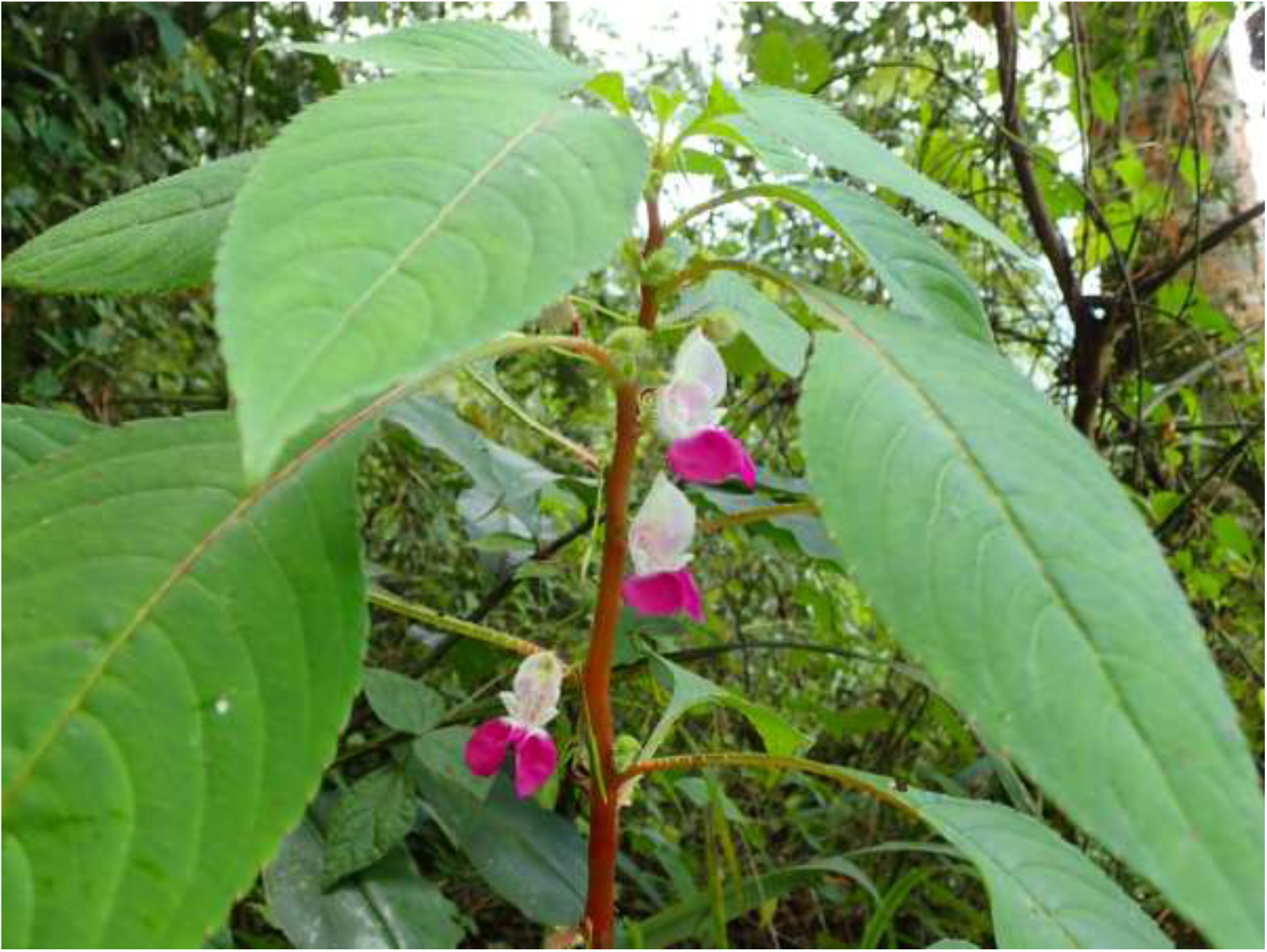
Impatiens banen. Habit, flowers in frontal view. *van der Burgt* 2373. Photo by Xander van der Burgt, Dec. 2019

**Fig. 2.**
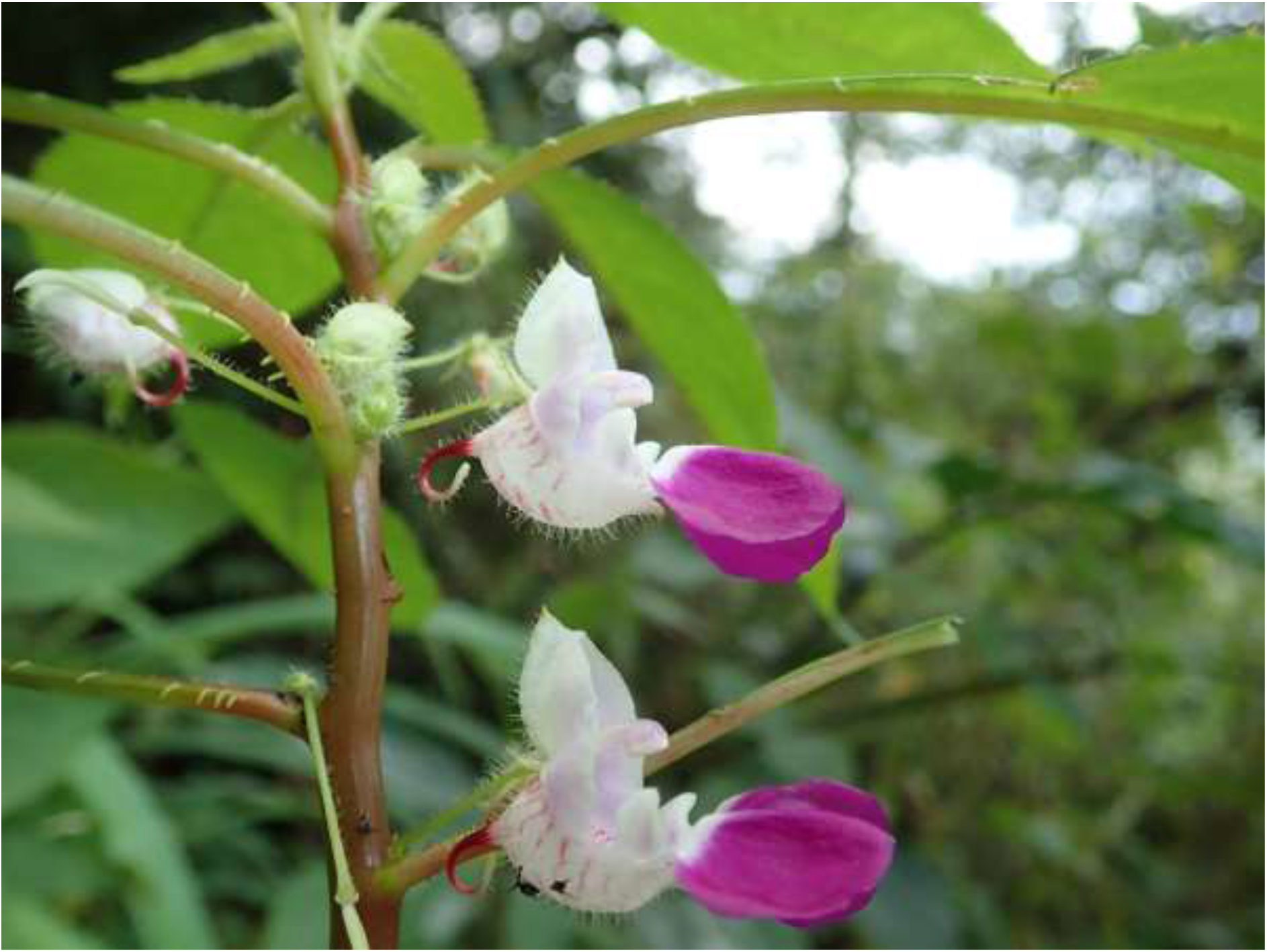
Impatiens banen. Close-up of flowers, side view. *van der Burgt* 2373. Photo by Xander van der Burgt, Dec. 2019

*Impatiens* L. with over 1067 species accepted (Plants of the World Online, continuously updated), are one of the most species-diverse genera of vascular plants, and are nearly cosmopolitan, indigenous species being absent only from S. America and Australia. The taxonomic framework for continental Africa was established by Grey-Wilson (1980) who recognised 110 species. Since then, 22 further species have been published (Abrahamczyk *et al*. 2016; Bos 1991; Cheek and Csiba 2002; Cheek and Fischer 1999; Fischer 1997; Fischer *et al*. 2003; 2021; Frimodt-Möller and Grey-Wilson 1999; Grey-Wilson 1981; Hallé and Louis 1989; Janssens *et al*. 2009a; 2010; 2011; 2015; 2018; Pócs 2007). The three broad centres of species diversity in tropical Africa are the western African mountains, primarily the Cameroon Highlands (28 species), the E. Arc mountains of Tanzania with the Kenya Highlands (24 species) and the Albertine Rift (20 species) (Fischer *et al*. 2021).

Africa was colonised by *Impatiens* from SW China on three occasions (Janssens *et al*. 2009b). The first colonisation was in the Early Miocene (clade A1, E. and S. Africa). Two further colonisations occurred in the Late Miocene or Early Pliocene, namely clade A2, endemic to W. Africa, and clade A3, which gave rise to the largest diversification in Africa, mostly in E and E-Central Africa, but with some species in both W and C Africa and then generally geographically disjunct between these two locations, e.g., *Impatiens mannii* Hook. f. and *I. burtonii* Hook. f. (Janssens *et al*. 2009b). Most speciation has occurred in the Pleistocene and has been rapid, e.g., two Gabonese species which are calculated to have diverged from their sister species only 0.18 million years BP (Janssens *et al*. 2011). Many of the species are geographically localised, several to individual mountains. In Africa, *Impatiens* characterise humid tropical and subtropical montane forests above 500 – 800 (− 5000) m alt. Most species of *Impatiens* cannot survive drought or extended exposure to direct sunlight. As a result, *Impatiens* species are typically confined to stream margins, waterside boulders, and wet and/or montane forests (Fischer, 2004). The species are pollinated by insects, including bees and butterflies, but with at least ten species which are adapted in multiple ways to being pollinated by sunbirds (Bartoš et al. 2012; Bartoš & Janeček 2014; Hořák & Janeček 2021). Studies of flower structures in relation to insect pollination currently lag behind those for bird pollinated species. Seed dispersal is mediated by explosive fruits (Grey-Wilson 1980).

## Materials & Methods

This study is based on herbarium specimens. All specimens cited have been seen unless indicated as “n.v.”. The methodology for the surveys in which most of the specimens were collected is given in Cheek & Cable (1997). Herbarium citations follow Index Herbariorum (Thiers *et al*. continuously updated), nomenclature follows Turland *et al*. (2018) and binomial authorities follow IPNI (continuously updated). The Flore du Cameroun volume for Balsaminaceae (Grey-Wilson 1981) followed by the monograph of African *Impatiens* (Grey-Wilson 1980) were the principal reference works used to determine the identifications of the specimens of what proved to be the new species. Material of the suspected new species was compared morphologically with protologues, reference herbarium specimens, including type material of W-C African *Impatiens* principally at K, but also using material and online images from BR, MO, P and YA.

Points were georeferenced using locality information from herbarium specimens. The conservation assessment was made using the categories and criteria of IUCN (2012). Herbarium material was examined with a Leica Wild M8 dissecting binocular microscope fitted with an eyepiece graticule measuring in units of 0.025 mm at maximum magnification. The drawing was made with the same equipment using a Leica 308700 camera lucida attachment.

## Taxonomic Results

### Impatiens banen

This species first came to the attention of the first author in January 2022 when reviewing photos publicly available on Digifolia [https://dams.kew.org] and searching on ‘Ebo Forest’. The seven images there of *van der Burgt* 2373 collected in December 2019, had been identified as *Impatiens burtonii* Hook. f., and *Impatiens banen* indeed bears many similarities to this species: the proportions and size of the flower are close to each other, the flower is predominantly white, the mouth is gaping, orbicular in frontal view. Moreover, the lower sepal and spur, the dorsal petal, pedicel are long-hairy, and the upper surface of the leaf-blade is also hairy. However, in the key to species in Flore du Cameroun (Grey-Wilson 1981:7) this specimen would key out at couplet 14, not as *Impatiens burtonii*, but as *I. mannii* Hook f. on balance, because the petioles are fimbriate at the base, and the lower sepal has transverse bands of purple or pink (however, it is not glabrous as in *I. mannii*). Dissection of the flowers shows further shared characters of *Impatiens banen* with *I. mannii*: both species have a long stipe to the united lateral petals (sessile in *I. burtonii*) and the upper of these united petals is elevated above the horizontal and protrudes from the mouth of the lower sepal in both species (vs. held below the horizontal in the same plane as the lower petal in *I. burtonii*). It seems likely that these two are sister species.

Searches for additional specimens showed that the species was first collected in 2006 (*Osborne* 85, K, YA identified as *Impatiens sp*.), and had also been collected in 2014 (*Droissart* 1664 BRLU n.v, MO n.v., YA n.v. images viewed on GBIF, identified as *Impatiens cf. burtonii*), both also from inselbergs at Ebo. Table 1 below shows the characters separating *Impatiens banen* from *I. burtonii* and *I.mannii*.

**Table 1.** Diagnostic characters separating *Impatiens banen* from *I. burtonii* and *I.mannii*. Characters for the last two species from Grey-Wilson (1980; 1981) and specimens at K.

|  | *Impatiens burtonii* | *Impatiens banen* | *Impatiens mannii* |
| --- | --- | --- | --- |
| Petiole base fimbriate | No | Yes | Yes |
| Indumentum of adaxial leaf-blade | White hairy appressed | White hairy appressed | Glabrous (except veins) |
| Pink-purple transverse bands on lower sepal | Absent | Conspicuous | Conspicuous |
| Indumentum of pedicels, lower sepal & dorsal petal | Patent white hairy | Patent white hairy | Glabrous |
| Upper of the lateral united petals elevated above the horizontal | No | Yes | Yes |
| Base of lateral united petals | Sessile | Stipitate | Stipitate |
| Colour of petals | Uniformly white sometimes tinged slightly pink | Bicolored, the lower of the united lateral petals vivid pink-purple, upper and dorsal petal white | Uniformly white, sometimes tinged slightly pink |
| Mouth of perianth, viewed from front | Orbicular | Orbicular | Laterally compressed |
| Spur curvature & colour | Straight or curved <90°, white or slightly pink | Curved gradually along its length through nearly 360°, purple with white apex | Straight or curved <90°, white or slightly pink |
| Dorsal posture | Cucullate (hooded), margins not reflexed | Cucullate (hooded), margins reflexed | ± flat, erect or slightly reflexed |
| Apex of upper of lateral united petals | Rounded | Rounded | Acute |

#### Impatiens banen

Cheek *sp. nov*. Type: Cameroon, Littoral Region, Yingui-Yabassi area, Ebo Forest, Ebo Forest Research Station - Bekongo trail - open expanse of rock at 245 m along the trail, fl. fr. 6 Oct. 2006, *Osborne* 85 with Bekokon, Enang Abwe, Beheng (holotype K000634597, isotype YA). (Fig. 1 – 3)

**Fig. 3.**
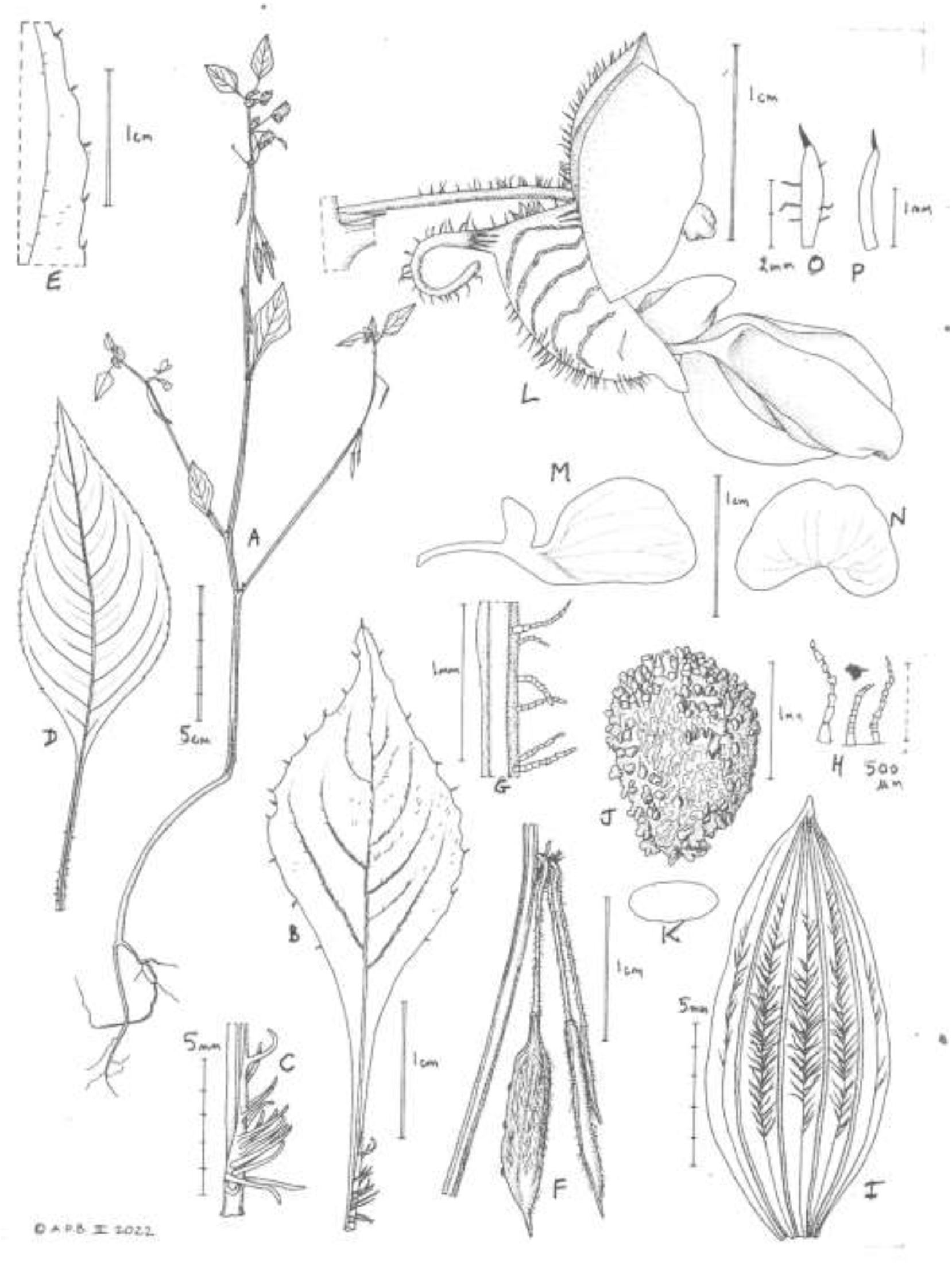
Impatiens banen. **A.** habit, flowering & fruiting plant; **B.** leaf-blade adaxial surface showing part of the surface indumentum and the basal fimbriae; **C.** detail of fimbriae in **B**; **D.** leaf-blade, showing large bladed variant; **E.** margin of leaf-blade showing teeth; **F.** fruit, showing geotropic habit; **G.** hairs on pedicel; **H.** multicellular hairs on fruit surface; **I.** fruit, opened flat, showing outer surface; **J.** seed, side view; **K.** outline of seed, transverse section; **L.** flower, side view; **M.** united lateral petals, showing long stipe; **N.** dorsal petal, pressed flat. **O.** lateral sepal; **P.** bract. **A-C, E-K, M-P** from *Osborne* 85; **D** & **L** from *van der Burgt* 2373. Drawn by ANDREW BROWN.

*Epilithic*, probably perennial herb 0.1 – 0.6 m tall; stems succulent, decumbent, rooting at the lower nodes, moderately branched, internodes 1.1 – 4(− 8) cm long, glabrous. *Leaves* spirally arranged, petiole (0.8 −)1.1 – 5.0(− 5.3) cm long, 1 – 1.8 mm wide, with numerous (4–7) fimbriae on each side along the edges, usually densely crowded together in the basal 5 – 7 mm, fimbriae cylindric-slightly curved, (1 −)1.5(− 2.25) x 0.2 – 0.4 mm, glabrous. *Lamina* ovate or lanceolate, 1.5 – 12.7(− 14.2) x (0.7 −)0.9 – 5.4(− 6.0) cm, apex acuminate, base cuneate-decurrent; lateral veins 6 – 8 on each side of the midrib, margin crenate to crenulate, adaxial surface uniformly covered in white, broad hairs, (0.5 −)0.75(− 0.9) mm. *Flowers* in contracted axillary racemes, appearing fasciculate, peduncles 0 – 1 mm long, rhachis contracted, 3 – 5-flowered, white, with pink-purple lower petals of the lateral united petals and pink-purple transverse bars on the lower sepal. *Bracts* numerous at raceme apex, linear-subulate (0.7 −)2–2.25 mm long, 0.1 – 0.3 mm, glabrous or with a few hairs. *Pedicels* suberect in flower, declinate (geotropic) and slightly accrescent in fruit, 11 – 15 mm long, hairs patent, multicellular, 0.5 – 0.7 mm long. *Lateral sepals* linear-lanceolate, 1.5 – 4 × 0.25 – 0.6 mm acute, nearly glabrous to densely long-hairy on the abaxial surface, hairs as pedicel. *Lower sepal* navicular, 7 – 13 mm long, 2.5 – 4 mm deep, terminating in a mucro 0.5 × 0.25 mm, abruptly constricted on one side into a dark red c. 10 mm long filiform spur gradually incurved along its length through 180 – 340 degrees, nearly forming a complete circle c. 3 – 4 mm diam., terminating in a bright white ellipsoid, 0.75 × 0.25 mm tip, spur white long-hairy throughout, hairs as pedicel, 0.5 – 0.7 mm long. *Dorsal petal* transversely reniform 9 – 15 × 10.5 – 12 mm when flattened, minutely apiculate, in life hooded (cucullate), erect, 6 mm wide, dorsally with a long crest 0.5 – 0.7 mm wide terminating in a mucro 0.3 × 0.3 mm, abaxial surface with 2.0 – 2.5 mm long, patent hairs along the veins of each side, hairs as pedicel, 1 mm long. *Lateral united petals* 20 – 22 mm long, on a long, slender stipe 7 – 8 × 0.8 – 1.5(− 2) mm, the upper petal much smaller than the lower; upper petal rectangular, rounded-quadrangular 10 – 11 × 8 – 10.5 mm apex rounded; lower petal concave, oblong 2.1 – 4 × 1.2 – 3.5 mm apex broadly rounded. *Androecium* white 4.7 – 8 mm long, staminal head 1.7 – 2 mm long. *Ovary* glabrous. *Fruit* fusiform, 14 – 16 × 1.2 – 3 mm apex acute, surface with 4 longitudinal pale bands; interband areas with a longitudinal line of spreading hairs, as pedicel, up to 1 mm long. *Seeds* ovoid, 1.5 × 1.1 mm, purple-brown the entire surface evenly scattered with dark gold conical projections each 0.12(− 0.25) × 0.075 mm.

##### RECOGNITION

*Impatiens banen* differs from *I. mannii* Hook. f. in that the mouth of the flower is orbicular (not laterally compressed) and the apex of the upper of the lateral united petals is rounded, not acute, the dorsal petal points forward and is hooded (not reflexed and flat), moreover the highly curved (through nearly 360°) purple spur with white tip is seen in neither *I. mannii, I. burtonii* nor any other Cameroonian species excepting *I. letouzeyi* Grey-Wilson where only the tip is curved (not the entire length of the spur). Above all *Impatiens banen* appears to be unique in Cameroon in the bicolored united lateral petals, the upper being white as in the rest of the flower, while the lower are an intense pink-purple. Additional diagnostic characters are given in table 1.

##### DISTRIBUTION

Cameroon, Littoral Region. So far only known from two inselbergs in the Ebo Forest.

##### SPECIMENS EXAMINED. CAMEROON

Littoral region, Ebo Forest Research Station - Bekongo Trail, fl.fr. 6 Oct. 2006, *Osborne* 85 with Bekokon, Enang Abwe, Beheng (holo. K000634597, iso. YA n.v.); ibid, reserve de faune d’Ebo, village de Ndokbaguengue. Campement de Djouma, sommet après le transect “Gachaka” Inselberg, ourlet herbacé, alt. 1003 m, fl. 15 Feb. 2014, *Droissart* 1664 with Couvreur & Kamdem 1664 (BRLU n.v., MO2971761, YA n.v.); ibid Ebo, Gashaka Hill 2 km northeast of Njuma Camp. 4° 21’ 29.4” N, 10° 14’ 59.4” E; fl. 3 Dec. 2019, *van der Burgt* 2373 (K001381835, MO, P, WAG, YA n.v.).

##### HABITAT

Restricted to open surfaces with seasonal seepages on granitic inselbergs, growing with seasonal wetland annual herbs such as species of *Scleria, Utricularia, Panicum, Selaginella* and mosses in lowland to submontane forest area; c. 850 m alt.

##### CONSERVATION STATUS

It is possible that *Impatiens banen* will yet be found at additional locations in Cameroon. However, while surveys have not been exhaustive, many thousands of specimens have been collected in areas to the north, south, west and east of Ebo, (Cheek 1992; Cheek *et al*. 1996; Cable & Cheek 1998; Cheek *et al*. 2000; Maisels *et al*. 2000, Chapman & Chapman 2001; Harvey *et al*. 2004; Cheek *et al*. 2004; Cheek *et al*. 2006; Cheek *et al*. 2010; Harvey *et al*. 2010; Cheek *et al*. 2011). If the species occurs elsewhere, that is most likely to be in at Mt Kupe in the Bakossi area to the NW since several range-restricted species are confined to these two locations, eg. *Coffea montekupensis* (Stoffelen *et al*. 1997). Moreover, several inselbergs, the habitat of *Impatiens banen*, occur in Bakossi (Cheek *et al*. 2004). Yet *Impatiens banen* is a flamboyant, long-flowering and immediately identifiable species so it would be surprising if it had been overlooked in such a well-studied area as Mt Kupe, suggesting that it is genuinely confined to the inselbergs in the forests of Ebo. Among the threats to inselbergs in forest in C. Africa reported in Pollard *et al*. (2006) are quarrying of the rock, increased frequency of fires e.g. from adjacent slash and burn agriculture, colonisation by non-indigenous species such as *Ananas comosus* L., which can out-compete indigenous inselberg species and replace them. Inselbergs are extensively targeted and quarried for their granite, which converted to gravel is in high demand for production of concrete and road surfacing and which can totally destroy the inselbergs concerned in terms of their plant communities (Couch *et al*. 2019).

*Impatiens banen* has an area of occupation of about 8 km^2^ using the 4 km^2^ cells requested by IUCN (2012). The extent of occurrence we estimate as about the same, using the IUCN approach. However, since this species appears specific to one particular inselberg microhabitat: sloping, open (and not tree or shrub covered) granitic inselberg surfaces with seepages, the area actually occupied is probably smaller. At the first known site in the eastern half of Ebo, near Bekob, Osborne, the first known collector of this species, estimates it as having been seen over an area of about 40 square metres only, in October 2006, with about 100 plants in total, and did not see it elsewhere. At the second known site in the western part of Ebo, van der Burgt observed hundreds of plants on the Gashaka or Gachaka inselberg in December 2019 and estimates that are potentially vastly more than that there are possibly 100 other inselbergs inside Ebo that have never been botanically explored. However, not all inselbergs necessarily have the specific inselberg micro-habitat in which the species seems restricted. The two known inselberg sites with *Impatiens banen* are within a single threat-based location in the sense of IUCN. The Ebo forest is not protected and was designated as a logging concession in Feb. 2021 (Lovell 2020) although this was suspended by the President of Cameroon in Aug. 2021 (Kew Science News 2020), the forest remains threatened by development as a future logging concession, for oil palm plantations and, for an open cast iron-ore mine (Cheek *et al*. 2018a). None of these, should they go ahead, will necessarily directly affect the inselbergs. However, they might do so indirectly if gravel is required, especially the last of which, if it goes ahead, will have requirements for this material for associated infrastructure such as a rail line. Therefore, for the present we here assess *Impatiens banen* as VU D2, but should the threatened iron ore mine go ahead the species will likely need to be reassessed as CR B1ab(iii) +B2ab(iii) (Critically Endangered).

##### ETYMOLOGY

Named as a noun in apposition for the Banen people, the guardians of the forest of Ebo, in Littoral Region, Yabassi.

##### PHENOLOGY

Flowering October to February.

##### NOTES

That *Impatiens banen* appears restricted to inselbergs in forest places it with several other range-restricted epilithic species restricted to this habitat in Cameroon and Gabon which have been revealed by the studies of Parmentier (2002; 2003; 2005; Parmentier & Mϋller 2006; Parmentier *et al*. 2006). These include *Coleus inselbergii* (B.J. Pollard & A.J. Paton) A.J. Paton (Pollard *et al*. 2006), *Oronesion testui* Raynal (Gentianaceae), *Polystachya odorata* Lindl. subsp. *gabonensis* Stévart (Orchidaceae), *Gladiolus mirus* Vaupel (Iridaceae) (Parmentier 2002, 2005) and *Cyperus inselbergensis* Lye (Cyperaceae, Lye 2013).

The inselbergs of Ebo remain to be surveyed, mapped and characterised. *Impatiens banen* is the first new species to science to be described from this habitat at Ebo. *Nothodissotis barteri* (Hook.f.) Ver.-Lib. & G.Kadereit (Melastomataceae) is an additional range-restricted and threatened (Vulnerable) species of the Ebo inselbergs, forming large shrubs to small spreading trees on domed surfaces (Veranso-Libalah *et al*. 2019).

*Impatiens banen* is a rare case of an apparently inselberg-specific *Impatiens.The* only other such species known to us is *Impatiens floretii* Hallé & A.M. Louis of Gabon (Hallé & Louis 1989). That inselberg-specific species are so rare in *Impatiens* is unexpected given that the genus as whole has a predilection for growing on damp rock surfaces (see introduction) so would appear to be pre-adapted for this habitat.

##### The Ebo Forest, Littoral Region

The Ebo Forest, a former proposed National Park, covers c. 1,400 km^2^ of lowland and submontane forest, with an altitudinal range of 130 – 1115 m alt. and a rainfall of 2.3 – 3.1 m p.a (Abwe & Morgan 2008; Cheek et al. 2018a). To date 84 globally threatened species of plant have been documented including 16 new to science, of which eight are globally endemic to Ebo: *Crateranthus cameroonensis* Cheek & Prance (Prance & Jongkind 2015), *Inversodicraea ebo* Cheek (Cheek *et al*. 2017), *Kupeantha ebo* M.G. Alvarez & Cheek (Cheek *et al*. 2018b), *Pseudohydrosme ebo* Cheek (Cheek *et al*. 2002a), *Talbotiella ebo* Mackinder & Wieringa (Mackinder *et al*. 2010), *Palisota ebo* Cheek (Cheek et al. 2018a), *Ardisia ebo* Cheek (Cheek & Xanthos 2012), while the most recent addition to the endemic species of Ebo was named *Uvariopsis dicaprio* Cheek & Gosline to honour Leonardo DiCaprio who championed conservation of Ebo in 2020 (Gosline *et al*. 2022).

*Impatiens banen* is the ninth published endemic plant species of Ebo and also increases to nine the number of taxa of *Impatiens* recorded from the 2950 specimens collected to date from Ebo, the remaining eight taxa being: *Impatiens filicornu* Hook. f., *Impatiens frithii* Cheek, *I. hians* var *hians* Hook.f., *I. kamerunensis* subsp. *kamerunensis* Warb., *I. mackeyena* subsp. *zenkeri* Hook. f., *I. mackeyana* subsp. *mackeyana* Hook. f., *I. macroptera* Hook. f. & *I. mannii*.

### Impatiens etugei

*Impatiens etugei* was first collected by Martin Etuge during a botanical reconnaissance of the Kom Wum Forest Reserve (*Etuge* 4759, K, YA 14 Nov. 2000) in NorthWest Region Cameroon. It was collected at a second site nearby in this forest nearly a year later (*Etuge* 4353, K, YA, 11 Oct. 2001) during a more extensive survey of plants in that forest, by a team of about 20 botanists and Earthwatch volunteers. The two Etuge specimens were tentatively identified “? *Impatiens letouzeyi”* by Iain Darbyshire in 2004, and later considered by him to be a possible new subspecies of *Impatiens mackeyana*. Placement in the *Impatiens macroptera* aggregate (Grey Wilson 1980) with these two species is indicated by the lowermost pair of teeth on the leaf blade margin being revolute and pointed at the leaf apex, by the lateral united petals which greatly exceed the saccate lower sepals and form a prominent, far-exserted lip. The Kom Wum material described here as *Impatiens etugei* differs from both the aforementioned species by the characters indicated in table 2 below.

**Table 2.**
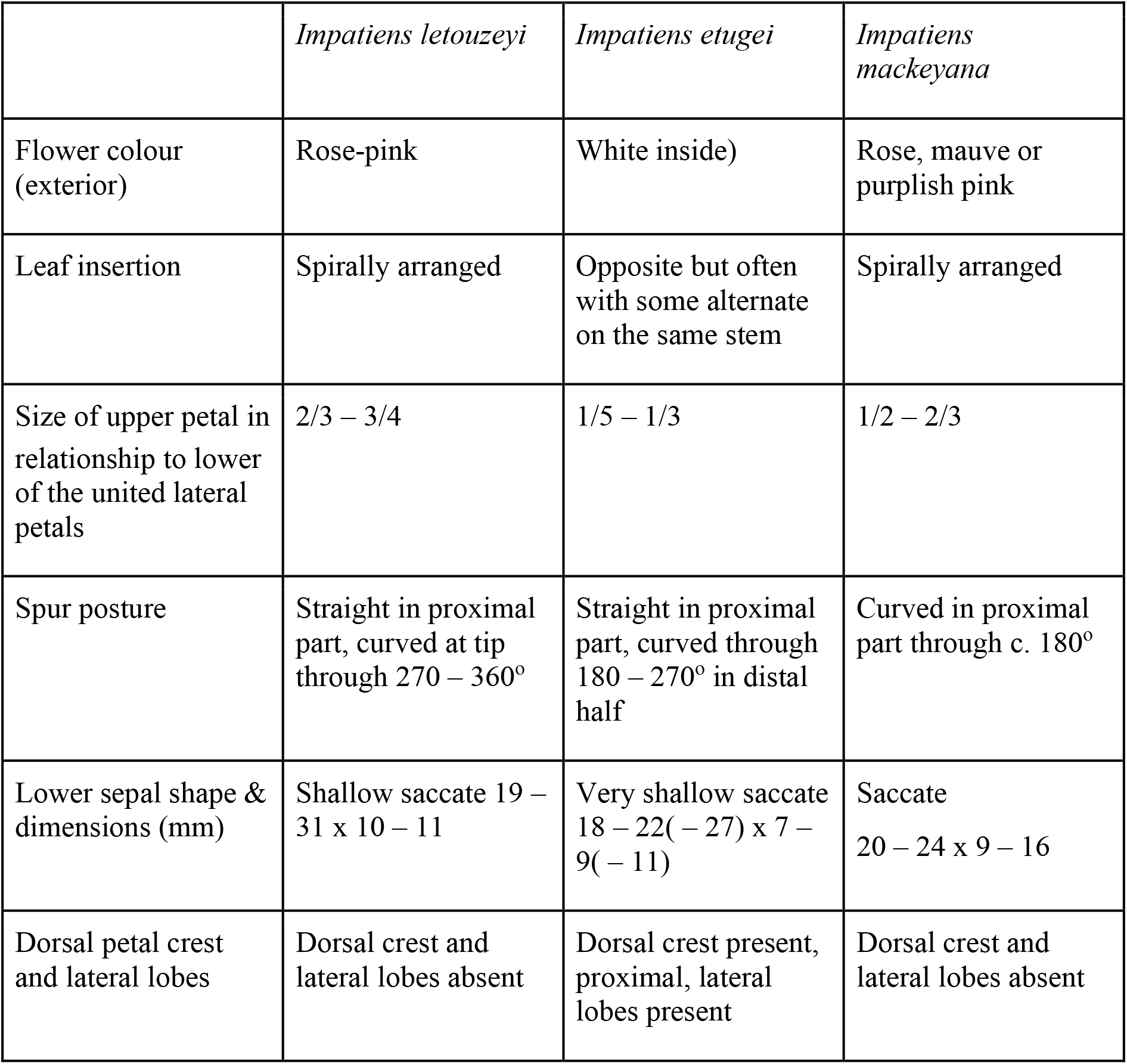
Diagnostic characters separating *Impatiens letouzeyi, Impatiens mackeyana* and *Impatiens etugei*. Data for the first two species from Grey Wilson (1980; 1981), Cheek *et al*. (2004) and specimens at K.

#### Impatiens etugei

Cheek sp. nov. Type: Cameroon, Northwest Region, Mencham Division, Bu, Nkom-Wum Forest Reserve, growing beside Mutel river on rocks 6° 16.50’N, 10° 06.70’ E, alt. 650 m, fl.fr. 14 Nov. 2000, *Etuge* 4759 with Gosline, Raza, Hugh, Bong, Fangha (holotype: K barcode K000593348, isotypes ETH, US, YA n.v.). (Fig. 4)

**Fig. 4.**
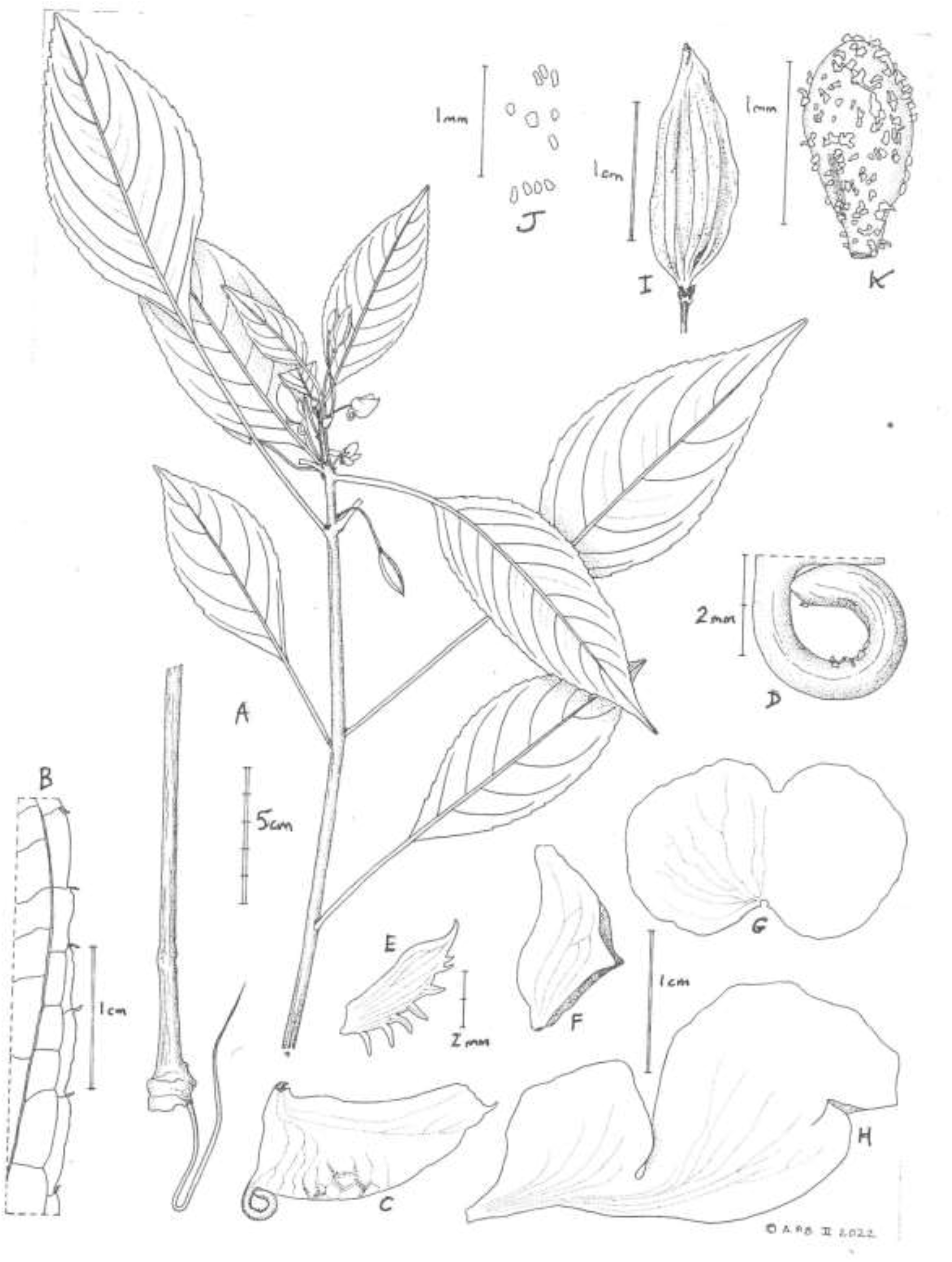
Impatiens etugei. **A.** habit, flowering & fruiting plant; **B.** leaf-blade abaxial surface portion showing margin; **C.** lower sepal and spur, side view; **D.** detail of spur from C; **E.** lateral sepal (hydrated); **F.** dorsal petal, side view showing lateral projections; **G.** dorsal petal, flattened; **H.** united lateral petals; **I.** fruit; **J.** fruit surface showing cystoliths; **K.** seed, side view. **A-E, G-J** from *Etuge* 4759; **F-K** from *Etuge* 4553r. Drawn by ANDREW BROWN.

*Epilithic, probably perennial herb* 0.15 – 1 m tall. *Stems* succulent, swollen, 5 – 9 mm diam. at base (dried specimens), leaves opposite or subopposite, often with some alternate on the same stem (Fig. 4A); internodes 0.3 – 1.6(− 5.8) cm long, glabrous or rarely (one plant) with erect brown simple hairs 0.1 – 0.25 mm long, moderately dense on stem, petioles, midribs and pedicels. *Leaves* elliptic, elliptic-oblong or lanceolate, (5.5 −)7 – 15.5(− 17) × (2.2 −)3 – 5.5(− 8) cm, obtuse or shortly acuminate, base acute-decurrent, secondary nerves (5 −)6 – 8(− 9) on each side of the midrib, margin serrate to crenate with patent fimbriae (Fig. 4B), the basal teeth pair strongly reflexed and directed towards the apex of the blade, junction with petiole with a pair of patent, red fimbriae 0.75 – 2 × 0.1 mm. Petioles (1 −)1.5 – 8.7(− 10.5) cm long, 0.2 cm wide, lacking fimbriae, glabrous. *Inflorescences* axillary, 1 – (2)-flowered, sessile. *Bracts* 5 – 8, rosulate, red, narrowly triangular to epedunculate linear, 1.5 – 2 × 0.2 mm, glabrous. *Pedicel* 20 – 24 × 1 mm, in fruit accrescent, 30 – 33 mm long. *Flowers* white on the outside, pink-orange or orange on the inside (probably the upper of the lateral united petals). *Lateral sepals* 2, ovate or lanceolate 5 – 6(− 8) × (2 −)2.5 – 3.5(− 4) mm, apex acute, terminating in a mucron, base subcordate, posterior margin entire, anterior margin with (4 −)5 – 7(− 8) teeth, the most proximal tooth longest (1 −)2 – 3 × 0.2 mm, reflexed, protruding above the corolla, teeth becoming shorter towards the sepal apex, midrib slightly raised, flanked with 2 white longitudinal nerves on each side of the midrib (Fig. 4E). *Lower sepal* shallow-saccate, slipper-like, 18 – 22(− 27) mm long, 7 – 9(− 11) mm deep, with c. 5 transverse purple bars, abruptly constricted into a 7 – 9(− 10) mm long filiform spur, proximal part straight, the distal part curved through 180 – 270°, apex slightly dilated, rounded (Fig. 4C). *Dorsal petal* cucullate, c. 14 × 21 mm when flattened, in plan view (pressed herbarium specimen Fig. 4G), basal sinus c. 3 × 6 mm, in lateral view 18 × 7 mm (Fig. 4F), apex acuminate-erect; dorsal crest c. 1.5 × 4 mm, proximal, lateral lobes c. 1 × 1 mm, apex rounded, inserted 8 mm from base. *Lateral united petals* 32 – 41 mm long; upper petal ⅕ – ⅓ the size of the lower petal of each pair, broadly oblong., or subtriangular, entire; lower petal oblong to dimidiate elliptic (14 −)18 – 25(− 28) × (10 −)12 – 15(− 18) mm, with a 3 mm deep emargination on the inner margin c. 6 mm from the apex (Fig. 4H). *Ovary* glabrous. *Fruit* elliptic-dimidiate, c. 1.7 × 1.1 cm apex acuminate-rostrate (Fig. 4I). *Seeds* narrowly ovate in the side view c. 1.1 × 0.75 mm flattened, 0.3 mm wide, red, with deep yellow longitudinal raphides c. 0.25 mm long, evenly curved, c. 50% of surface, with flat, mucilaginous white hairs 0.05 – 0.2 mm long (Fig. 4K).

##### RECOGNITION

*Impatiens etugei* is closely similar to *I. letouzeyi* Grey-Wilson and *I. mackeyana* Hook. f., resembling the first in the spur which is straight in the proximal half, curving in the distal part (vs curved from base in *I. mackeyana*). It resembles the second in the narrow lateral sepals which are toothed on one side only (vs. broad and toothed on both sides in *I. letouzeyi*). It differs from both species in that the leaves are usually opposite and subopposite, as well as alternate, on one stem, the dorsal petal has lateral projections from the margin (vs always alternate and lateral projections absent) and the flower exterior is white (vs shades of pink or pink-purple).

##### DISTRIBUTION

Cameroon, Northwest Region, Mencham Division. So far only known from the Nkom-Wum Forest Reserve near Bu village.

##### SPECIMENS EXAMINED. CAMEROON

North West Region (formerly Province), Mencham Division, Bamenda, Nkom-Wum Forest Reserve, growing beside Mutel river on rocks 6° 16.50’N, 10° 06.70’ E, alt. 650 m, fl.fr. 14 Nov. 2000, *Etuge* 4759 with Gosline, Raza, Hugh, Bong, Fangha (holotype: K barcode K000593348, isotypes ETH, US, YA n.v.). Bu-Sasey forest valley, 668 m alt., forest along Meteh River, fl. 11 Oct. 2001, *Etuge* 4353r with Ndong Emmanuel (K000593347, G, MA, YA n.v.).

##### HABITAT

On rocks in river in lowland evergreen forest; 650 – 670 m alt.

##### CONSERVATION STATUS

*Impatiens etugei* has an area of occupancy of 8 km^2^ and extent of occurrence estimated to be of the same area, using the 4 km^2^ cells preferred by IUCN. This taxon has not been found in surrounding areas, despite the numerous surveys cited above under *Impatiens ebo*. Threats to the Kom Wum forest are recorded in detail by Lyong (2020) including incursions by farmers from local communities clearing forest for crop fields, pasture and farm buildings, illegal logging, logs and planks being floated down streams from the reserve for sale in neighbouring Nigeria. These are threats to *Impatiens etugei* because removal of forest shade from *Impatiens* can be deleterious to their survival (see introduction), and log floating can scrape clean river boulders of plant life. Two specimens (cited above) from what is thought to be the single threat-based location are recorded. Therefore we assess this taxon as Critically Endangered, CR B1+B2a,b(iii).

##### ETYMOLOGY

Named in honour of the late Martin Etuge Ekwoge (-2020) of Bakossi, known professionally as Martin Etuge, botanical specimen collector from 1984 – 2000, mainly in Southwest and Northwest Regions of Cameroon, initially with Duncan W. Thomas (MO), based at Kumba. He was employed as a horticulturist at the Limbe Botanic Garden, Mount Cameroon Project c. 1991 – 1995, then became based at Nyasoso as botanical consultant with the San Diego Zoo’s former Mount Kupe Conservation Project, also working with e.g. RBG, Kew through the Earthwatch programme as a freelance botanical collector. Duplicates of his many herbarium specimens, numbering about 5000, can be found in herbaria around the world. He is also commemorated by the taxa *Cola etugei* Cheek (Cheek *et al*. 2020a), *Kupea martinetugei* Cheek & S.A. Williams (Cheek *et al*. 2003) and *Psychotria martinetugei* Cheek (Cheek & Csiba 2002b).

##### NOTES

Both specimens of *Impatiens etugei* are comprised of five sheets, derived probably from as many plants, they are from two different sites. The two subpopulations differ from each other vegetatively. The leaf-blades of *Etuge* 4759 are larger, and more slender in proportion to their length, and on longer petioles, compared with those of *Etuge* 4353r. The plants of all the sheets of both collections are completely glabrous, except one plant (one sheet) of *Etuge* 4759 which is moderately densely brown patent hairy on the distal two internodes, petioles, abaxial midribs and pedicels.

The seeds of *Impatiens etugei* appear to have mucilaginous hairs. One seed was found attached to a leaf-blade. Therefore, to test for adhesivity, five seeds from a fruit were rehydrated, applied to other surfaces such as paper, allowed to dry and found to adhere. The long white flat papillae-like projections from the seedcoat become swollen, erect and colourless when hydrated. We hypothesise that seeds might be dispersed from one rock in a river to another, carried on the bodies of primates such as chimpanzees (zoochory). In areas of intact forest habitat with low human impact, large rocks in rivers are often frequented by primates as evidenced by deposits of faeces (Cheek pers. obs. 1992-2014), and Chefor Fotang has recorded this for chimpanzees in Kom Wum (pers. comm. to Cheek, February 2022).

##### The Kom Wum Forest Reserve

*Impatiens etugei* is only known from the Kom Wum Forest Reserve, also known as Nkom Wum. This is one of the few surviving intact patches of evergreen lowland to submontane forest in Northwest Region, Cameroon. The area of the Kom Wum Forest Reserve is c. 80 km^2^ over an altitudinal range of 565 – 1640 m alt. and with a mean annual rainfall of 2.4 m p.a, the wet season extending from mid-March to mid-October. Kom Wum is home to seven diurnal and six nocturnal primate species (Fotang *et al*. 2021). The plant species of Kom Wum are imperfectly known and there is no checklist of species yet available. *Impatiens etugei* is probably the first species of plant to be described from the reserve, however, a second species, *Uvariopsis etugei* Dagallier & Couvreur (Annonaceae) is in the course of publication (Couvreur *et al*. in press), which is known from Kom Wum and only one other location outside. Therefore, it is likely that additional range-restricted and globally threatened plant species will be found at Kom Wum so long as natural habitat remains there. Since December 2016 access to the forest and all of SouthWest and NorthWest Regions has been restricted due to the armed conflict between the secessionists and national government.

## Conclusions

Until species are delimited and known to science, it is much more difficult to assess them for their conservation status and so the possibility of protecting them is reduced (Cheek *et al*. 2020b). The majority of new species described today tend to be range-restricted, making them especially likely to be threatened, although there are some exceptions (e.g. Cheek & Etuge 2009; Cheek *et al*. 2019a). 133 new vascular plant species to science have been published from Cameroon in the last five years (IPNI, continuously updated)

To maximise the survival prospects of range-restricted species there is an urgent need to formally assess the species for their extinction risk, applying the criteria of a recognised system, of which the IUCN Red List of Threatened Species is the most widely accepted (Bachman *et al*. 2019). The majority of plant species still lack such assessments (Nic Lughadha *et al*. 2020). Documented extinctions of plant species are increasing (Humphreys *et al*. 2019) and recent estimates suggest that as many as two fifths of the world’s plant species are now threatened with extinction (Nic Lughadha *et al*. 2020). In Cameroon *Oxygyne triandra* Schltr. is globally extinct as is *Afrothismia pachyantha* Schltr. (Cheek & Williams 1999; Cheek *et al*. 2018c; Cheek *et al*. 2019b). In some cases, species appear to be extinct even before they are named for science, such as *Vepris bali* Cheek (Cheek *et al*. 2018d), and in neighbouring Gabon, *Pseudohydrosme bogneri* Cheek & Moxon-Holt (Moxon-Holt & Cheek 2021). Most of the >800 Cameroonian species in the Red Data Book for the plants of Cameroon are threatened with extinction due to habitat clearance or degradation, especially of forest for large-scale plantations e.g. oil palm and small-holder agriculture, following logging (Onana & Cheek 2011). Efforts are now being made to delimit the highest priority areas in Cameroon for plant conservation as Tropical Important Plant Areas (TIPAs) using the revised IPA criteria set out in Darbyshire *et al*. (2017).

National governments and leaders have recognised the importance of species assessed as threatened on the Red List and documented in IPAs or TIPAs as demonstrated recently in Cameroon when in part due to the high number of plant species on the Red List (Lovell 2020), a logging concession was revoked for the Ebo forest (Kew Science News 2020).

It is hoped that formal publication and Red Listing of additional threatened endemic species such as the *Impatiens banen* and *I. etugei* will help support the motivation and case for resources for the protection of the natural areas in which they occur, the Ebo and Kom Wum forests respectively.

## Acknowledgements

Completion of work on the Cameroon species in this paper was through support of the Cameroon TIPAs programme from Players of People’s Postcode Lottery (PPL).

Led by Ekwoge Abwe and Bethan Morgan, The Ebo Forest Research Programme supported by San Diego Zoo Wildlife Alliance have been crucial in inviting and helping us to access that wonderful forest with its Banen and Bassa communities to discover and support the protection of threatened plant species unknown to science such as *Impatiens banen*.

John De Marco and Anne Gardner of the Bamenda Highlands Forest Project based at Anajuya, North West Region, together with the late Paul Mzeka of the Apiculture and Nature Conservation Organisation (ANCO) of Bamenda, encouraged our survey of the Nkom Wum Reserve which resulted in the collection of *Impatiens etugei*. Sincere thanks to them all. We also thank Chefor Fotang for recent discussion on the area of Kom Wum.

The specimens cited of *Impatiens etugei* in this paper were collected with the support of volunteers and sponsored scientists arranged by Earthwatch Europe, Oxford.

Drs Satabié, Achoundong and more recently Florence Ngo Ngwe,, Jean Betti Lagarde, the current and former directors, of IRAD (Institute of Research in Agronomic Development)-National Herbarium of Cameroon,, and their staff are thanked for expediting the collaboration between our two institutes under the terms of our Memorandum of Collaboration. Janis Shillito typed the manuscript. Two anonymous reviewers and Xander van der Burgt are thanked for constructively reviewing an earlier version of this paper.

The authors declare no conflicts of interest.

